# The Geometry of Partial Fitness Orders and an Efficient Method for Detecting Genetic Interactions

**DOI:** 10.1101/180976

**Authors:** Caitlin Lienkaemper, Lisa Lamberti, James Drain, Niko Beerenwinkel, Alex Gavryushkin

## Abstract

We present an efficient computational approach for detecting genetic interactions from fitness comparison data together with a geometric interpretation using polyhedral cones associated to partial orderings. Genetic interactions are defined by linear forms with integer coefficients in the fitness variables assigned to genotypes. These forms generalize several popular approaches to study interactions, including Fourier-Walsh coefficients, interaction coordinates, and circuits. We assume that fitness measurements come with high uncertainty or are even unavailable, as is the case for many empirical studies, and derive interactions only from comparisons of genotypes with respect to their fitness, i.e. from partial fitness orders. We present a characterization of the class of partial fitness orders that imply interactions, using a graph-theoretic approach. Our characterization then yields an efficient algorithm for testing the condition when certain genetic interactions, such as sign epistasis, are implied. This provides an exponential improvement of the best previously known method. We also present a geometric interpretation of our characterization, which provides the basis for statistical analysis of partial fitness orders and genetic interactions.

## 1. Introduction

Genetic interactions - or dependence of the fitness effect produced by a set of mutations on the genetic background - play an important role in determining evolutionary trajectories of populations. For instance, a set of individually beneficial mutations may exhibit diminishing returns: while the combined effect of all mutations is beneficial, it is not as beneficial as one would expect given the individual mutations, for example the sum of all single mutation effects. Diminishing returns have been shown to slow down the pace of adaptation (Chou et al. 2011). Similarly, the combined effect of a set of mutations can be stronger than the effect expected from the individual mutations. This so-called synergistic epistasis between deleterious mutations has been observed for instance in human populations, where it affects the distribution of deleterious alleles in the genome (Sohail et al. 2017). Genetic interactions can affect not just the magnitude of the effect of a mutation, but also the sign of the effect. That is, a particular mutation may have a beneficial effect in one genetic background and a deleterious effect in another. This type of interaction, often termed sign epistasis, can act to constrain evolutionary trajectories by requiring that some mutations occur before others (Gong, Suchard, and Bloom 2013; Kvitek and Sherlock 2011; Weinreich, Watson, and Chao 2005).

In this work, we are concerned with detecting the presence of genetic interactions from limited genotype-phenotype data. We understand an interaction as a deviation from a null linear model. For example, the two-way interaction is defined as a deviation from the null assumption that

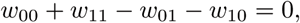

and the total three-way interaction as a deviation from the null assumption that

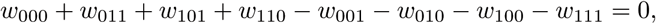

where the binary indices correspond to biallelic genotypes (e.g. 00 is the wild type and 11 is the double mutant), and w denotes fitness (see Section 2 for detailed definitions). Empirical data sets produced in interaction studies typically provide replicate measurements of a trait, such as antibiotic or other forms of resistance, for various genotypes. The trait is often chosen to measure fitness, or approximations thereof, and used to infer genetic interactions in the fitness landscape. However, in many cases the association between the trait and fitness is unclear. For example, comparative fitness assays (or competition experiments) produce data uninformative with respect to the distribution of the absolute fitness values of genotypes. Furthermore, assuming only comparative genotype-phenotype data, one detects the sign but not the magnitude of the interactions. This makes standard statistical methods such as ANOVA or the multiple regression framework inapplicable.

Here, we make a conservative assumption that the only available information about fitness is the (partial) ranking of genotypes, that is, we assume that no precise fitness measurements of all genotypes are available, a frequent situation in practice, for example, due to measurement noise. Specifically, we assume either that some of the information in a total order of all fitness values must be discarded, or that the only signal available in the data is of the form “genotype *g*_1_ has higher fitness than genotype *g*_2_”, for various pairs of genotypes. We assume that these pairwise fitness comparisons are consistent with each other, i.e., the relation they define is transitive. We call this type of data a *partial fitness order*, which is a partial order of genotypes with respect to their fitness. Data of this type arises for example by considering certain directed acyclic graphs where the vertices are given by genotypes and where the edges between genotypes are directed towards the genotype with higher fitness. Importantly, our assumptions make our methods widely applicable as the same techniques can be applied to other types of rankings and not just genotypes ranked with respect to their fitness. For example, a partial fitness order of genotypes can be obtained in studies that involve measuring various kinds of breeding values (see for example Habier, Fernando, and Dekkers 2007) or other applications of generalised linear models.

Even though we assume that accurate fitness measurements cannot be accessed for all genotypes in a given genotype space, our methods still enable us to detect interactions implied by the partial fitness order. In particular, we are able to detect sign epistasis. Moreover, even when a complete fitness ranking of genotypes is available a method to detect interactions from a partial order alone might provide additional insight. Indeed, there are cases where the rank order method fails to reveal interactions even though interactions in the system can be detected from the actual fitness measurements (Crona, Gavryushkin, et al. 2017). In this case, partial fitness order methods can be used to conclude diminishing returns or synergistic epistasis and refute sign epistasis. In addition, if one is only interested in certain fitness comparisons (for instance, comparisons between mutational neighbors), our methods yield a criterion to determine whether these comparisons are sufficient to imply interaction. These observations will have implications for the evolutionary trajectories of the system and provide insight as to the mechanism of the interaction (Weinreich, Watson, and Chao 2005). A detailed analysis of the class of linear orders that imply interactions, including relevant literature and data, can be found in (Crona, Gavryushkin, et al. 2017).

In this paper, we build on the results obtained by Crona, Gavryushkin, et al. (2017) and settle the most general case of ranking data, when only a partial fitness order is available. Specifically, we present a graph-theoretic characterization of the class of partial fitness orders that imply genetic interactions (Section 3), thus establishing the conjecture presented by Gavryushkin at the Interactions between Algebra and the Sciences conference held in Max Planck Institute for Mathematics in the Sciences, Leipzig, on 27 May 2017. Second, we use our characterization to derive a cubic time algorithm for testing the condition when a given partial order implies such an interaction (Section 3). This provides an exponential improvement of the best previously known method that involves iterating through the list of all possible linear extensions of the given partial order (Crona, Gavryushkin, et al. 2017). Third, we count all partial orders for up to 8 labeled elements which imply an interaction (Section 4), and fourth, we provide a geometric interpretation of our characterization, useful for statistical analysis of partial fitness orders that imply genetic interactions (Section 5). Finally, we use public data sets to demonstrate our methods (Section 6) and conclude by describing possible future directions for this line of research (Section 7).

## 2. Technical introduction

In this section we introduce all necessary terminology and notations and describe a basic statistical approach to probabilistic inference of (higher-order) interactions from partial fitness orders, which further motivates our results on partial fitness orders.

Consider a system *𝒢* consisting of *k* genotypes, where *k* is a positive integer. We denote the fitness of a genotype *g* ∊ *𝒢* by *w*_*g*_, which is a real number. Typically, we take fitness to be a positive real number, though this does not affect our arguments. All fitness measurements (*w*_*g*_)_*g∊𝒢*_ together then determine what is called the *fitness landscape* associated to *𝒢*. We refer to the tuple of all the fitness measurements defining a fitness landscape by *W* = (*w*_*g*_)_*g∊𝒢*_.

We define genetic interactions using linear forms with integer coefficients in the following way. Let *ƒ* be a linear form in variables *W* with integer coefficients which sum to 0, that is,

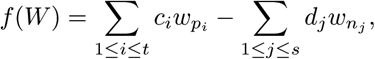

such that

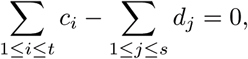

where all *c*_*i*_, *d*_*j*_, *t*, and *s* are positive integer numbers such that *t* + *s* = *k* and *𝒢* = {*p*_1_,…,*p*_*t*_, *n*_1_,…, *n*_*s*_}. We then say that the fitness landscape *has positive ƒ-interaction* if *ƒ*(*W*) > 0. When *ƒ*(*W*) < 0, we say that *has negative ƒ-interaction*. Hence *ƒ*-interactions are a special case of what is known in applied statistics as contrasts. From now on, we will focus on determining positive *ƒ*-interactions unless we explicitly state otherwise. Everything in this paper extends straightforwardly to the negative case. This definition of *ƒ*-interaction generalizes several well-known and widely used notions of gene interaction, including Fourier-Walsh coefficients, interaction coordinates, and circuit interactions (Weinreich, Lan, et al. 2013; Beerenwinkel, Pachter, and Sturmfels 2007). The detailed exposition including illustrative examples comparing all these approaches can be found in (Crona, Gavryushkin, et al. 2017, Materials and methods).

Estimating the probability of *ƒ*-interaction in an empirical setting amounts to quantifying the uncertainty of *ƒ*(*W*) > 0 given some comparative fitness data. Although in this setting the fitness values *w*_*g*_ are unavailable, the comparative fitness data typically allows to deduce a partial fitness order. For example, this type of data could arise from replicate measurements of any quantitative trait monotonic with respect to fitness. Another example of such data is fitness comparison data produced in competition or survival experiments, which are a popular method for fitness estimation.

Sometimes a partial fitness order is enough to deduce *ƒ*-interactions and hence estimate the probability of interaction. We use the following definition to address such situations. We say that a partial fitness order 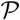 = (*𝒢*, ≺) *implies positive ƒ-interaction* if *ƒ*(*W*) > 0 whenever *W* satisfy the partial order 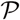 that is, *w*_*g*_ < *w*_*h*_ for all *g ≺ h*

Then the probability of *ƒ*-interaction given comparative fitness data can be estimated by considering the probability support of all partial orders that imply *ƒ*-interaction. However, such an estimation would require the condition of whether or not a partial order 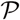 implies *ƒ*-interaction to be checked routinely for different partial orders 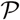. Hence the complexity of this condition might become a computational bottleneck in practice. In this paper we resolve this problem by designing an efficient polynomial-time algorithm for checking whether or not a partial order implies *ƒ*-interaction.

We note that the fact that a partial order does not imply positive or negative *ƒ*-interaction does not necessarily mean that the system does not have *ƒ*-interaction. This situation therefore should be interpreted as the partial order being non-informative with respect to the *ƒ*-interaction; the issue is discussed in more detail in (Crona, Gavryushkin, et al. 2017).

We now review the results from (Crona, Gavryushkin, et al. 2017) about how to detect whether a rank order implies *ƒ*-interaction. In the terminology used in our paper, this situation corresponds to the case when the partial fitness order is a linear order (also called a total order or a rank order). We then study arbitrary partial orders, and present an efficient algorithm for determining when a partial order implies *ƒ*-interaction.

For a linear form *ƒ* as above, define a function *ϕ*^*ƒ*^ from the set of (strict) linear orders on the set of genotypes to words over the alphabet {*P, N*} as follows. The linear order on *k* genotypes

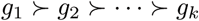

is mapped to a word *ω* consisting of *c*_*i*_ copies of *P* if *w*_*g*_*i*__ has a positive coefficient *c*_*i*_ in *ƒ*; and *d*_*i*_ copies of *N* if *w*_*g*_*i*__ has a negative coefficient *d*_*i*_ in *ƒ*. The subwords of *ω* corresponding to different *g*_*i*_’s appear according to the linear order (from highest to lowest). See (Crona, Gavryushkin, et al. 2017) for various examples of this construction.

The following result from (Crona, Gavryushkin, et al. 2017) depends upon the characterization of linear orders which imply interaction in terms of Dyck words.

### Definition 1.

Let Σ be an alphabet consisting of two letters *P* and *N*. A *Dyck word* is a word *ω* consisting of an equal number of *P*’s and *N*’s such that every prefix of *ω* contains at least as many *P*’s as *N*’s.

For example, *PPNNPN* and *PNPNPN* are Dyck words, but *PPNNNP* is not, since the prefix *PPNNN* contains more *N*’s than *P*’s. Clearly the definition of Dyck word does not depend on the choice of alphabet one considers. Dyck words arise in a number of contexts in discrete mathematics. For instance, they are in bijection with full binary trees with *n* + 1 leaves. If the symbols *P* and *N* are replaced with “(” and “)”, a string is a Dyck word only if all parenthesis are correctly matched. For more on Dyck words, see (Stanley 2001).

### Theorem 1

(Crona, Gavryushkin, et al. (2017)). *A linear order* 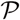 = (*𝒢*, ≺) *on the set of genotypes implies positive ƒ-interaction if and only if* 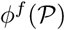 *is a Dyck word that starts with P*.

The statement of this theorem can equivalently be formulated by saying that a linear order 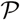 implies positive *ƒ*-interaction if and only if there exists a partition of the set of all genotypes *𝒢* into pairs (*p*_*i*_, *n*_*j*_) such that *p*_*i*_ *≻ n*_*j*_ for all *i,j*, where each *p*_*i*_ appears in *c*_*i*_ pairs and each *n*_*j*_ in *d*_*j*_ pairs (see Crona, Gavryushkin, et al. 2017). As we will see in the next section, this formulation can be generalized to arbitrary partial orders. Note that here and also below we slightly abuse notation because for certain choices of *c*_*i*_’s and *d*_*j*_’s, we do not have a partition in the strict sense, but this technical difficulty can be avoided by, for example, distinguishing between the copies of *p*_*i*_ and *n*_*j*_.

## 3. Efficient method to infer interactions from partial orders

In this section we generalize Theorem 1 and characterize partial orders which imply *ƒ*-interactions. We then apply our characterization to design an efficient algorithm for testing the condition of whether or not a partial order implies *ƒ*-interaction.

### Theorem 2.

*Let 𝒢* = (*p*_1_,…,*p*_*t*_,*n*_1_,…,*n*_*s*_) *be a set of genotypes*,

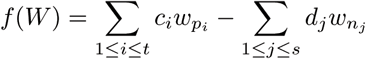

*be a linear form, and* 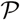 = (*𝒢*, ≺) *be a partial order on 𝒢. Then* 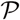 = (*𝒢*, ≺) *implies positive ƒ-interaction if and only if there exists a partition of the set of all genotypes 𝒢 into pairs* (*p*_*i*_, *n*_*j*_), *p*_*i*_ ≻ *n*_*j*_ *where each genotype p*_*i*_ *appears in exactly c*_*i*_ *pairs and each genotype n*_*j*_ *in exactly d*_*j*_ *pairs*.

Before we proceed with the proof, let us illustrate Theorem 2 with the following example. Let

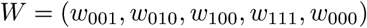

and consider the linear form:

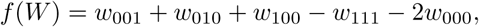

which measures the deviation of the triple mutant’s fitness from its expectation based on single mutants only, and corresponds to one of the circuit interactions characterized in (Beerenwinkel, Pachter, and Sturmfels 2007). We see that there is positive *ƒ*-interaction implied by the order of the fitness values if and only if for each fitness value with a positive coefficient in *ƒ*, there is a strictly smaller fitness value with negative coefficient. This smaller fitness value must be unique, unless the absolute value of the coefficient is greater than one, as in this case for *w*_000_.

*Proof*. The “if” implication is clear and follows from (Crona, Gavryushkin, et al. 2017). We provide here some details for convenience of the reader.

Recall that

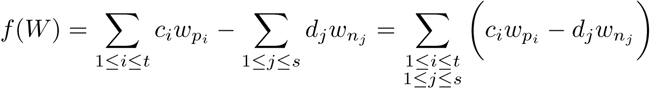

and if there is a partition for the partial order 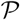 as claimed and the indices *i, j* are arranged according to the partition, each difference in the last sum is strictly greater than 0, thus *ƒ*(*W*) > 0.

In order to prove the converse statement, we prove its contrapositive. That is, we prove that if there is no such partition of the set of all genotypes, *ƒ*-interaction is not implied.

Construct the bipartite graph *G* = (**P ⊔ N**, *E*), where **P** contains *c*_*i*_ copies of *p*_*i*_ for each *i* and **N** contains *d*_*j*_ copies of *n*_*j*_ for each *j*, such that (*n*_*j*_,*p*_*i*_) ∈ *E* whenever *n*_*j*_ ≺ *p*_*i*_ in the partial order 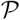. The number of elements in **P** and **N** coincide, since the coefficients of *ƒ* sum to zero. Observe that a partition of the set of all genotypes *𝒢* into pairs (*n*_*j*_, *p*_*i*_) such that *n*_*j*_ ≺ *p*_*i*_ for all *i, j* as in the theorem is exactly a perfect matching in *G*. By Hall’s theorem, there exists a perfect matching in *G* if and only if for each *S* ⊆ N, we have

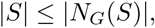

where *N*_*G*_(*S*) ⊆ **P** is the set of vertices in *G* adjacent to a vertex in *S* (Diestel 2016). Suppose that there is no partition of the genotypes into pairs as above, and thus no perfect matching in *G*. Then there is some subset *S* ⊆ **N** such that

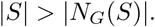

Using this subset *S*, we construct a linear extension of the partial order 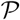 which does not map to a Dyck word under the map *ϕ*^*ƒ*^ introduced above.

To do so, we first reindex N so that

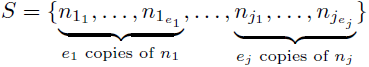

and the order *n*_1_ ≻…≻ *n*_*j*_ is compatible with 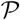. Note that *d*_1_ ≥ *e*_1_ ≥ 1,…, *d*_j_ ≥ *e*_j_ ≥ 1.

Let 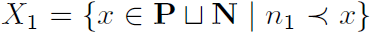 be the set of elements in 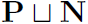 ordered above *n*_1_. Note that 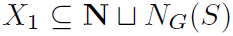, since if 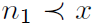 and 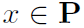, then by construction, 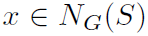. Consider a linear extension 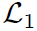 of the partial order 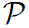 on *X*_1_ given by 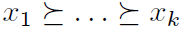. Extend 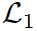 with all duplicates 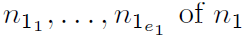. That is, consider the linear extension 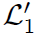 given by

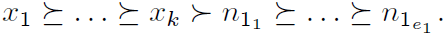

Now, we repeat this process with *n*_2_ and the rest of elements in *S*. Let *X*_2_ be the set of elements in **P** ⊔ **N** which are required by 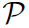 to be strictly greater than *n*_2_ but are not contained among the elements included in 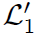. Again, 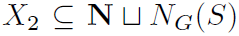. Extend the previous linear order 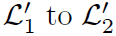 as follows, maintaining the property that 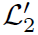 is compatible with 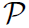:

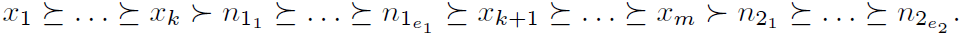

We repeat this process for the rest of elements in S in the same way. To complete the construction, we observe that every remaining element (not included in 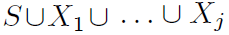 is compatible with being smaller than all elements of 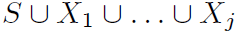. Hence, we can sort all remaining elements and add them at the end of the last linear order 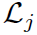 as smaller elements, to obtain a linear order 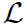 which extends 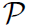.

The linear order 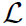 does not map to a Dyck word (starting in *P*) under 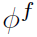, since the image of the smallest prefix of 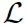 which contains all elements of S contains more *N*’s than *P*’s since 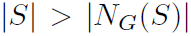. Therefore, by Theorem 3 from (Crona, Gavryushkin, et al. 2017), this partial order does not imply positive *f*-interaction.

A variant of Theorem 2 was independently obtained in (Crona and Luo 2017).

We now proceed by designing an efficient algorithm for detecting whether or not a partial fitness order 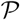 implies *ƒ*-interaction. Recall that *ƒ*(*W*) = Σ_1≤*i*≤*t*_*c_i_w_p_i__* - Σ_1≤*j*≤*s*_*d_j_w_n_j__* and Σ_1≤*i*≤*t*_*c_i_* = Σ_1≤*j*≤*s*_*d_j_*. In the next result, denote m = Σ_1≤*i*≤*t*_*c_i_*.

### Theorem 3.

*There exists an 𝒪*(*m*^3^) *algorithm to determine whether a partial order* 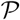 = (*𝒢*, ≺) *implies positive *ƒ*-interaction.*

*Proof.* We use the notation introduced in the proof of Theorem 2. That theorem implies that a partial order implies that *ƒ*(*W*) > 0 if and only if there exists a partition of the set *P‘* ∪ *N‘* into pairs (*p_i_, n_j_*) such that *p_i_* ≺ *n_j_*. As noted in the proof of Theorem 2, this condition is equivalent to the existence of a perfect matching in the bipartite graph *G* = (**P, N**, *E*) such that (*n_j_, p_i_*) ∈ *E* whenever *p_i_* ≺ *n_j_*. Thus, we can detect whether the poset implies *ƒ*(*W*) > 0 by checking whether there exists a perfect matching in *G*.

Using the Hopcroft-Karp algorithm (Hopcroft and Karp 1971), we can find a perfect matching in a bipartite graph with *m* vertices in time *𝒪*(*m*^5/2^). Assume we are presented the partially ordered set 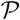 as a directed acyclic graph. Then we can construct the bipartite graph *G* in time *𝒪*(*m*^3^) using the following algorithm: for each *p* ∈ **P**, use breadth first search to find all elements of **N** which are reachable along a directed path from *p*, and add these to *p*’s list of neighbors. The worst-case complexity of this step is *𝒪*(*m*^2^). Repeat this process for each of the *m* vertices to obtain a representation of *G* as an adjacency list - that is, as a list containing a list of neighboring vertices for each vertex. Thus, since we can construct the graph in *𝒪*(*m*^3^) time and determine whether a matching exists in the graph in *𝒪*(*m*^5/2^) time, and *m*^3^ > *m*^5/2^, we can determine whether the partial order determines the sign of a linear form in *𝒪*(*m*^3^) time.

The complexity of our algorithm for computing whether a partial order on a set of genotypes implies *ƒ*-interaction depends not just on the number of genotypes, but also on the coefficients of *ƒ*. Specifically, the worst case complexity is a function not just of the number of genotypes, but also of the number of positive summands of *ƒ* (with multiplicity) Σ_1≤*i*≤*t*_*c*_*i*_. In most practical cases (circuits, interaction coordinates, etc.), however, this last number is small relative to the number of genotypes.

The two upper level steps of our algorithm are

- constructing the bipartite graph and
- detecting whether a perfect matching exists.

We think that the upper complexity bound *𝒪*(*m*_3_) achieved in this paper can be improved. In particular, an appropriate choice of data structures can reduce the complexity of constructing the bipartite graph, which is the most demanding step in our algorithm. However, this improvement goes beyond the scope of this work, as we found the algorithm’s performance sufficient on typical data sets used in higher-order interaction studies (see Crona, Gavryushkin, et al. 2017, for a survey of such data sets). See https://github.com/gavruskin/fitlands/tree/posets/posets.ipynb, where our implementation of the algorithm designed in Theorem 3 can be accessed.

Linear programming provides an alternate method to check whether a partial order implies interaction: checking whether a system is consistent with positive interaction amounts to checking that the feasible region of the linear program with constraints coming from the partial order and the constraint *ƒ* > 0 is nonempty. In this case, the complexity of the computation will just depend on the number of genotypes, not on the coefficients, thus with large coefficients, linear programming is a more practical approach then the one presented here. However, by providing a purely combinatorial description of when a partial order implies interaction, our method may be more useful for studying classes of partial orders.

## 4. Counting partial orders that imply interaction

In this section, we determine the proportion of partial orders on *k* = 2, 4, 6, and 8 genotypes that imply positive *ƒ*-interaction for an arbitrary fixed linear form *ƒ* with *k* coefficients 1 and −1 which sum to 0. Our results are summarized in Table 1.

**Table 1.** For *k* ∈ {2,4,6,8}, the number of partial orders implying positive *ƒ*-interaction, the number of partial orders not implying *ƒ*-interaction, and the proportion of partial orders implying either positive or negative *ƒ*-interaction, truncated to the second digit. Note that the number of partial orders implying negative *ƒ*-interaction is, by symmetry, equal to the number of partial orders implying positive *ƒ*-interaction. Hence, the proportion is obtained by dividing twice the number of partial orders implying positive interaction by the total number of partial orders.

| $k$ | positive interactions | no implication | proportion implying interaction |
| --- | --- | --- | --- |
| 2 | 1 | 1 | 0.67 |
| 4 | 31 | 157 | 0.28 |
| 6 | 10,876 | 108,271 | 0.17 |
| 8 | 22,217,743 | 387,287,893 | 0.10 |

We carried out our computations in the open-source mathematics software system Sage (http://www.sagemath.org) using the source code available at https://github.com/gavruskin/fitlands/tree/posets/. To make the computations possible, we employed the automorphism groups (Stein 2008) of our partial orders in the following way.

### Definition 2.

An *automorphism* of a partial order 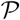 = (*V*, ≺) is a bijection *σ: V* → *V* such that if *u* ≺ *v*, then *σ*(*u*) ≺ *σ*(*v*). The set of automorphisms of a finite partial order forms a group, denoted by Aut(*G*), under the operation of composition.

In our enumeration, we observe that the number of different ways to label a poset with *k* elements is 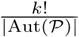. To see this, take a labeling of the elements on the poset. A permutation of the labels is an automorphism if and only if it does not change the partial order. Thus, each relabeling of the poset elements falls into an equivalence class of |Aut(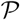)| relabelings that do not change the partial order. Thus, since there are *k*! ways to relabel the elements, there are 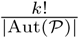 non-equivalent ways to label the poset.

As an example, consider the partial order

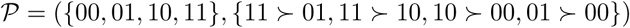

with Hasse diagram depicted on the left of the following figure:

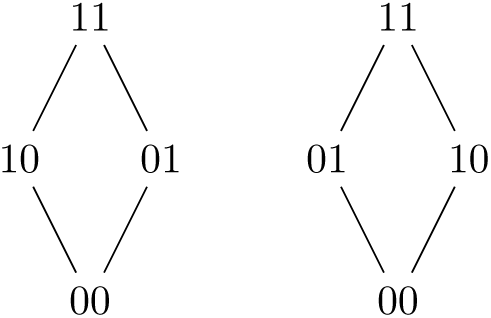

The relabeling to the Hasse diagram on the right yields the same partial order. Thus, this permutation switching 10 and 01 is an automorphism of the graph. However, any other permutation of the labels changes the partial order. Thus, this poset has an automorphism group of cardinality 2. Then since there are 4! = 24 ways to relabel the elements of {00,01,10,11}, there are 4!/2 = 12 different labelings of this poset.

The implementation of our algorithm proceeds as follows: Using a built in function in Sage, we produced a list of partial orders on *k* elements up to isomorphism. Since whether a partial order implies *ƒ*-interaction depends on the element labels, not just the structure of the partial order, we have to distinguish between different labeled partial orders within each partial order automorphism class. To achieve this, for each partial order (up to automorphism), we check for *ƒ*-interaction with each partition of the labels used by Sage into two classes of equal size. We let one class correspond to elements with coefficient 1 and one class correspond to elements with coefficient −1. Then each choice of partition corresponds to ((*k*/2)!)^2^ possible reassignments of the labels, since we can permute the elements within the class with coefficient 1 and the class with coefficient −1 without changing the partition. We compute the cardinality of the automorphism group for each partial order, and add 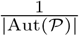 to our count of partial orders that imply positive, negative, or no *ƒ*-interaction respectively.

We note that as we increase *k*, the proportion of partial orders implying *ƒ*-interaction decreases. The sums of the values we obtained for the number of partial orders implying positive *ƒ*-interaction, the number of partial orders implying negative *ƒ*-interaction, and the number of partial orders not implying any *ƒ*-interaction appears in OEIS (*The On-Line Encyclopedia of Integer Sequences* 2017) as the number of partial orders on *k* labeled elements, as expected. However, none of the columns of our table appear in OEIS. Further, each number is either prime or has a prime factorization including fairly large primes. Thus, the number of partial orders, the number of partial orders implying *ƒ*-interaction, and the number of partial orders not implying *ƒ*-interaction do not appear to be given by any simple formula.

The final computation, for *k* = 8, took approximately two hours to complete on the SageMath Cloud server. The main factor affecting the performance of these computations is that the number of partial orders on *k* elements is super-exponential in the number of elements (*The On-Line Encyclopedia of Integer Sequences* 2017). Therefore, we did not count the number of partial orders on *k* elements which imply *ƒ*-interaction for *k* ≥ 10. Since we can still quickly check whether individual partial orders on *k* ≥ 10 elements imply *ƒ*-interaction, sampling methods can be used to estimate how the proportion of partial orders which imply interaction changes as we increase *k.*

Our computational observation that the fraction of partial orders which imply *ƒ*-interaction approaches 0 as *k* increases is true in general:

### Theorem 4.

*As k approaches infinity, the fraction of partial orders on which imply f-interaction approaches zero*.

*Proof.* Partition the set of partial orders on *k* elements into its isomorphism classes. We show that, within each isomorphism class, the proportion of partial orders implying positive *ƒ*-interaction is bounded by 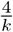 whenever *ƒ* is a linear form with coefficients ±1. Consider an arbitrary isomorphism class, and take an arbitrary labeled poset 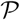 in this class. There are 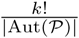 distinct labeled posets in this isomorphism class. There are 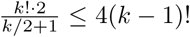 linear orders on *k* elements which imply positive *ƒ*-interaction (Crona, Gavryushkin, et al. 2017, Proposition 1). Now, take an arbitrary linear extension of 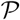. Each permutation of labels which takes 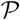 to a labeled poset which implies positive *ƒ*-interaction must take this linear extension to one that implies interaction, thus there are at most 4(*k - 1*)! permutations of labels which take our labeled poset to one that implies f interaction. Further, all |Aut(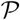)| permutations of labels which give the same labeled poset must take our linear order to one that implies interaction. Thus, there are at most 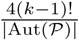 labeled posets in this isomorphism class that imply interaction. Thus, the proportion of posets in this isomorphism class that imply interaction is at most 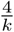.

## 5. Geometric interpretation

In this section, we present a geometric interpretation of Theorem 2, and derive some geometric, combinatorial, and statistical results. We denote the space ℝ^|*𝒢*|^ where |*𝒢*| is the number of elements in *𝒢* simply by ℝ^*𝒢*^ and index the coordinate axes by the unknown fitness values *w_g_*’s, where *g* ∈ *𝒢*. When we assume fitness values to be nonnegative, we work in the positive orthants of ℝ^*𝒢*^, which we denote by 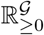. In this case, every point of 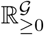 corresponds to a possible measurement of a fitness value for each genotype. When we do not assume fitness values to be nonnegative, every point in ℝ^*𝒢*^ corresponds to a fitness value for each genotype. The geometric interpretation of our results about partial orders uses the language of polyhedral cones.

### Definition 3.

A *convex polyhedral cone* in ℝ^*k*^ is a subset of ℝ^*k*^ cut out by a finite number of linear inequalities.

Like all cones, convex polyhedral cones are closed under the operations of taking nonnegative linear combinations of their elements. For the remainder of this paper, we will refer to convex polyhedral cones simply as cones. Let 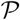 = (*𝒢*, ≺) be a partial order (which may be a total order). Then the set of points *x* ∈ ℝ_*𝒢*_ whose coordinate order satisfies 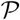 defines a cone. To see this, note that this set of points is cut out by a finite number of linear inequalities defined by the partial order. Hence this set is a convex polyhedral cone by definition. We call a cone associated to a partial order in this way an *order cone* and denote it by *C*_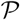_. Further, a linear form *ƒ* in the variables *w_g_*’s defines a hyperplane through the origin, where *ƒ* takes the value 0. The linear form is positive in the half space on one side of this hyperplane and negative on the other. Let *H*_*ƒ*,+_ be the half space on which f is positive, *H*_*ƒ*_ be the hyperplane on which *ƒ* is zero, and *H*_*ƒ*,-_ be the half space on which *ƒ* is negative. Then Theorem 2 has the following geometric interpretation. Recall that *ƒ*(*W*) = Σ_1≤*i*≤*t*_*c_i_w_p_i__* - Σ_1≤*j*≤*s*_*d_j_w_n_j__* is a linear form with integer coefficients that sum to zero, that is Σ_1≤*i*≤*t*_*c_i_* = Σ_1≤*j*≤*s*_*d_j_*.

### Theorem 5.

*The cone C*_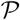_ ⊂ *H*_*ƒ*,+_ *if and only if there exists a partition of the set of all genotypes 𝒢 into pairs* (*p_i_, n_j_*) *such that p_i_* ≻ *n_j_ for all i, j, where each p_i_ appears in c_i_ pairs and each n_j_ in d_j_ pairs*.

*Proof.* Note that 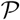 implies positive *ƒ*-interaction if and only if *C*_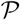_ ⊂ *H*_*ƒ*,+_. Then the claim follows from Theorem 2.

As a corollary of Theorem 5, we obtain the following result, which will be applied in our analysis of a Malaria data set in Subsection 6.

### Definition 4.

Let 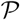_1_ and 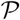*2* be partial orders. Their *intersection* is the partial order 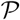_1_ ∩ 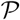_2_ such that *x* ≺_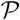_1_∩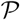_2__ *y* if and only if *x* ≺_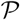_1__ *y* and *x* ≺_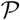_2__ *y*.

To illustrate the above definition consider the partial orders 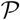_1_ = 00 ≻ 10 ≻ 01 and 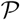_2_ = 10 ≻ 01 ≻ 00. Then 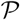_1_ ∩ 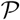_2_ is the partial order given by 10 ≻ 01.

### Corollary 1.

*Let **A** and **B** be linear orders such that C_**A**_, C_**B**_* ⊂ *H*_*ƒ*,+_, *then the order cone C_**A**⊂**B**_ of the intersection of **A** and **B** is contained in H*_*ƒ*,+_ *if and only if **A**∩**B*** = (*𝒢*, ≺) *satisfies the condition in Theorem 5.*

*Proof.* Immediately follows from Theorem 5.

Now, we prove the following theorem which is important for statistical analysis of interactions from fitness comparison data, for example, in the analysis of a sample from the probability distribution over the space of partial fitness orders with the aim to quantify the uncertainty of interactions. Specifically, we show the following. If two uncertain fitness measurements give distinct linear orders ***A*** and ***B*** both of which imply positive *ƒ*-interaction, then there exist fitness measurements corresponding to a sequence of linear orders which are intermediate between ***A*** and ***B*** in the sense that every linear order in this sequence is different from its predecessor by just one transposition of a pair of adjacent elements, and no linear order in the sequence implies negative *ƒ*-interaction.

### Theorem 6.

*Let U* ⊂ *ℝ*^*k*^ *be a path connected, open set which has a nonempty intersection with the cones C*_*A*_ *and C*_*B*_, *where* ***A*** *and* ***B*** *are linear orders on k elements. Then there exist linear orders 𝓛*_1_ = ***A***,…, *𝓛*_*n*_ = ***B*** *such that 𝓛*_*i*_ *and 𝓛*_*i*+1_ *differ by one adjacent transposition and U ∩ C*_*𝓛*_*i*__ ≠ *Φ for each 1 ≤ i < n*.

*Proof.* We denote cones *C*_*𝓛*_i__ by simply *C*_*i*_, for all *i*, throughout the proof. Note that the cones *C*_*i*_ and *C*_*j*_ have a (*k* − 1)-dimensional face as their intersection if and only if *𝓛*_*i*_ and *𝓛*_*j*_ differ by a single adjacent transposition. Further, note that any path which passes from *C*_*i*_ to *C*_*j*_ without passing through the interior of any other cone must pass through the intersection of the boundaries of *C*_*i*_ and *C*_*j*_. Either this boundary is an (*k* − 1)-dimensional face, and *𝓛*_*i*_ and *𝓛*_*j*_ differ by an adjacent transposition, or we pass through a lower dimensional face. In the latter case, this means a neighborhood of the point where we pass through the boundary contains all cones that intersect at this point. Thus, a neighborhood of this point contains a sequence of order cones of linear orders which differ by one adjacent transposition each.

Now, consider a path from a point in *U∩C*_*A*_ to a point in *U∩C*_*B*_. Since *U* is open and path connected, it contains some path of this form, as well as a neighborhood around the path. Consider the sequence of cones this path passes through - by the observation above, we know that if the path passes from *C*_*i*_to *C*_*j*_, either *𝓛*_*i*_ and *𝓛*_*j*_ differ by an adjacent transposition, or a neighborhood of the point where we pass from *C*_*i*_to *C*_*j*_ intersects the cones of a sequence of linear orders which differ by one adjacent transposition each. Thus, in either case, a path from *C*_*A*_ to *C*_*B*_ has a neighborhood that intersects the order cones of a sequence of linear orders *𝓛*_1_,…, *𝓛*_*n*_ such that *𝓛*_1_ = ***A***, *𝓛*_*n*_ = ***B***, and *𝓛*_*i*_ and *𝓛*_*i*+1_ differ by one adjacent transposition for each 1 ≤ *i* < *n.* These linear orders satisfy the claim of the theorem.

We illustrate Theorem 6 in Figure 1, where we depicted six regions corresponding to six slices of order cones in 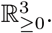 Each cone is associated to a linear order of three elements, *x* = *w*_*g*_*i*__, *y* = *w*_*g*_*j*__, *z* = *w*_*g*_*k*__. Points inside the cones correspond to different fitness values satisfying the linear order defining the given cone. Thus, for example a point inside *C*_*x>y>z*_ specifies three fitness values associated to three genotypes *g*_*i*_, *g*_*j*_, *g*_*k*_ ∈ *𝒢* and such that *w*_*g*_*i*__ > *w*_*g*_j__ > *w*_*g*_*k*__.

As a corollary to Theorem 6, we obtain the following result.

### Corollary 2.

*Suppose **A** and **B** are linear orders which imply positive *ƒ*-interaction. Then there exists a sequence of linear orders 𝓛*_1_ = *A,…,𝓛*_*k*_ = *B such that 𝓛*_*i*_ *and 𝓛*_*i*+1_ *for* 1 ≤ *i* < *n all differ by one adjacent transposition and no 𝓛*_*i*_ *implies negative ƒ-interaction*.

*Proof*. The half-space *H*_*ƒ*, +_ is a connected open set which has a nonempty intersection with *C*_*A*_ and *C*_*B*_. Thus, this result follows from Theorem 5. □

Spaces with complicated geometries and combinatorics are known to cause significant difficulties for statistical analysis (Billera, Holmes, and Vogtmann 2001). For example, the space of trees, which is a particular instance of the space of partial orders, required deep mathematical advances to understand basic statistics, such as confidence regions and convex hulls, over the space (Billera, Holmes, and Vogtmann 2001; Gavryushkin and Drummond 2016). Advances in the geometry and combinatorics of such spaces are the stepping stone for efficient statistical methods such as Markov Chain Monte Carlo (Gavryushkin, Whidden, and Matsen 2017; Dinh et al. 2017). In a similar vein, we expect that the approach of this section allows to efficiently study probability distributions over the space of partial fitness orders. Hence our results provide a theoretical foundation for statistical analysis of partial fitness orders.

**Figure 1.**
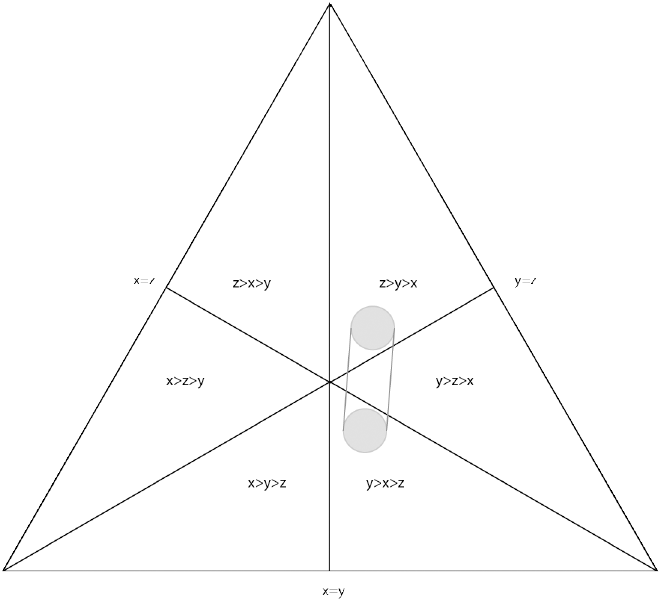
We show a slice of 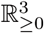, demonstrating that 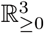 is divided into six cones corresponding to the six possible orders of the elements *x, y, z*. The convex hull of a ball in the order cone *C*_*z>y>x*_ and a ball in the order cone *C*_*y>x>z*_ passes through the order cone *C*_*y>z>x*_. Note that the order *y > z > x* differs by one adjacent transposition from both *z > y > x* and *y > x > z*.

## 6. Applications

In this section we illustrate how our results can be used in fitness-based genetic interactions studies.

### Malaria

As a first application, we consider the following three linear fitness orders inferred and analyzed by Ogbunugafor and Hartl (2016) (see also Crona, Gavryushkin, et al. 2017). These fitness orders have been obtained by measuring the growth rate of the parasite Plasmodium vivax exposed to various concentrations of the antimalarial drug pyrimethamine.

> *𝓛*_1_: 111 ≻ 011 ≻ 001 ≻ 101 ≻ 010 ≻ 100 ≻ 110 ≻ 000
>
> *𝓛*_2_: 111 ≻ 011 ≻ 001 ≻ 010 ≻ 100 ≻ 101 ≻ 110 ≻ 000
>
> *𝓛*_3_: 111 ≻ 011 ≻ 010 ≻ 001 ≻ 100 ≻ 110 ≻ 101 ≻ 000

As noted in (Crona, Gavryushkin, et al. 2017), each of these linear orders implies negative total 3-way interaction, which in our terms is *ƒ*-interaction where

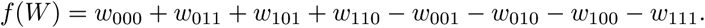

We now strengthen this conclusion using the approach developed in this paper. To do so, we consider only the shared rankings of the genotypes across different drug concentrations, that is, the partial fitness order 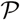 obtained by taking the intersection *𝓛*_1_ ∩ *𝓛*_2_ ∩ *𝓛*_3_. This intersection has the following Hasse diagram:

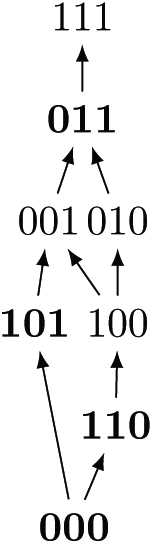

We then apply Theorem 2 and conclude that the obtained partial order 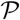 implies negative *ƒ*-interaction. Indeed, the three bottom genotypes with even number of ones (depicted in bold) can be matched to the three genotypes with even number of ones above them, and matching 011 to 111 completes the construction of a perfect matching in this graph.

This means that even if the only comparisons we are confident about are those that agree across the three drug concentrations, we can still conclude that this system exhibits negative total 3-way interaction. In particular, this approach shows that we can ignore the inconsistency of the three rank orders associated to the three different drug concentrations under inspection. Hence any tuple of fitness values W which satisfies every fitness comparison found in each linear order will imply interaction. Further, in this case our results from Section 5 imply that any point in the convex hull of the points we have measured will imply negative 3-way interaction. This region allows to incorporate the uncertainties of the measurements and provides a region within which those measurements can vary without breaking the conclusion of interaction.

Moreover, in this situation we observe that *𝓛*_2_ can be obtained from *𝓛*_1_ by two permutations (see Corollary 2), with the intermediate liner order also implying negative total 3-way interaction:

> *𝓛*_1_: 111 ≻ 011 ≻ 001 ≻ 101 ≻ 010 ≻ 100 ≻ 110 ≻ 000
>
> *𝓛*_2_: 111 ≻ 011 ≻ 001 ≻ 010 ≻ 101 ≻ 100 ≻ 110 ≻ 000
>
> *𝓛*_3_: 111 ≻ 011 ≻ 001 ≻ 010 ≻ 100 ≻ 101 ≻ 110 ≻ 000

The exact same argument applies to *𝓛*_2_ and *𝓛*_3_. In general, intermediate linear orders of this type are examples of sequences of linear orders between two given linear orders as described in Corollary 2. These intermediate linear orders, *𝓛*_*i*_, can then be used to suggest further fitness comparisons to be tested to validated the conclusion of negative *ƒ*-interaction in this data set.

To conclude this section we also observe that unlike the intersection *𝓛*_1_ ∩ *𝓛*_2_ ∩ *𝓛*_3_ of the three linear orders given above, the intersection of multiple linear orders, each implying *ƒ*-interaction, does not necessarily imply *ƒ*-interaction. To illustrate this situation, consider for example the following linear orders:

> *𝓛*_4_: 111 ≻ 011 ≻ 001 ≻ 101 ≻ 010 ≻ 100 ≻ 110 ≻ 000
>
> *𝓛*_5_: 111 ≻ 011 ≻ 010 ≻ 101 ≻ 001 ≻ 100 ≻ 110 ≻ 000

Both linear orders *𝓛*_4_ and *𝓛*_5_ imply negative total 3-way interaction, as each linear order maps to the Dyck word *NPNPNNPP*. However, the partial order *𝓛*_4_ ∩ *𝓛*_5_ = 111 ≻ 011 ≻ 001,010,101 ≻ 100 ≻ 110 ≻ 000 does not imply total 3-way interaction, since there is no perfect matching and *𝓛*_4_ ∩ *𝓛*_5_ extends to the following linear order:

> *𝓛*_6_: 111 ≻ 011 ≻ 101 ≻ 010 ≻ 001 ≻ 100 ≻ 110 ≻ 000;

which maps to a non-Dyck word *NPPNNNPP*.This examples thus shows that the property “implies *ƒ*-interaction” is not closed under the operation of order intersection and illustrates Corollary 1.

### TEM β-lactamase

As a second application, we consider the data produced in the antibiotic resistance study of the TEM-family of *β*-lactamase (Mira et al. 2015). In the table below we display the average growth rates of 16 genotypes grown in the antibiotic AMP in 12 replicates. The 16 genotypes include the wild type and all combinations of amino acid substitutions in TEM-50.

By using these growth rate as a measure of fitness, we deduce that this system has positive total 4-way interaction by directly computing:

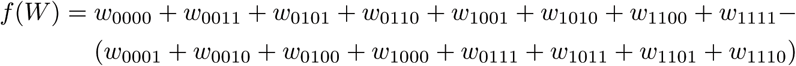

and noting that *ƒ*(*W*) > 0 for the values in Table 2.

**Table 2.**
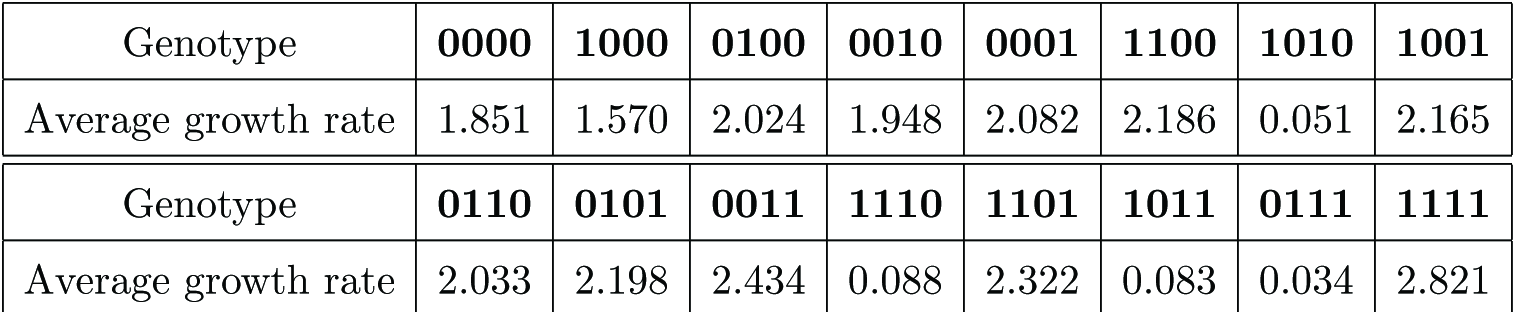
Average growth rates of the 16 genotypes grown in the antibiotic AMP.

| Genotype | 0000 | 1000 | 0100 | 0010 | 0001 | 1100 | 1010 | 1001 |
| --- | --- | --- | --- | --- | --- | --- | --- | --- |
| Average growth rate | 1.851 | 1.570 | 2.024 | 1.948 | 2.082 | 2.186 | 0.051 | 2.165 |
| Genotype | 0110 | 0101 | 0011 | 1110 | 1101 | 1011 | 0111 | 1111 |
| Average growth rate | 2.033 | 2.198 | 2.434 | 0.088 | 2.322 | 0.083 | 0.034 | 2.821 |

In the following, we demonstrate how our approach can be used to analyze the uncertainty of the conclusion that the system has positive total 4-way interaction. We start by ranking the genotypes according to their growth rates. This yields the linear order described in the following table (see Table 5 in (Mira et al. 2015)): In the following we assume that the average growth rates of these 16 genotypes vary but that they always preserve the ranking listed in Table 3. This assumption implies that the average growth rates map to the word

> *ω* = *PPNPPPNPNNPNNNPN*

according to the mapping described in Section 2. Reading *ω* from left to right, we notice that *ω* is a Dyck word. Hence, Theorem 1 can be used to confirm that the system has positive interaction. However, comparing the ranking and the average growth rates yields the following observation. The difference between the average growth rates assigned to the genotypes 0101, 1100, and 1001 as well as to the pairs of genotypes 0110, 0100 and 1110, 1011 is smaller comparing to the differences between all other average growth rates assigned to the remaining genotypes. See Figure 2 for a visual summary of this observation. These small differences in the growth rates highlight the difficulty of finding accurate and robust linear orders among the genotypes.

**Table 3.** Ranking of the 16 genotypes grown in the antibiotic AMP according to their average growth rates listed in Table 2.

| Genotype | 0000 | 1000 | 0100 | 0010 | 0001 | 1100 | 1010 | 1001 |
| --- | --- | --- | --- | --- | --- | --- | --- | --- |
| Rank | 11 | 12 | 9 | 10 | 7 | 5 | 15 | 6 |

| Genotype | 0110 | 0101 | 0011 | 1110 | 1101 | 1011 | 0111 | 1111 |
| --- | --- | --- | --- | --- | --- | --- | --- | --- |
| Rank | 8 | 4 | 2 | 13 | 3 | 14 | 16 | 1 |

**Figure 2.**
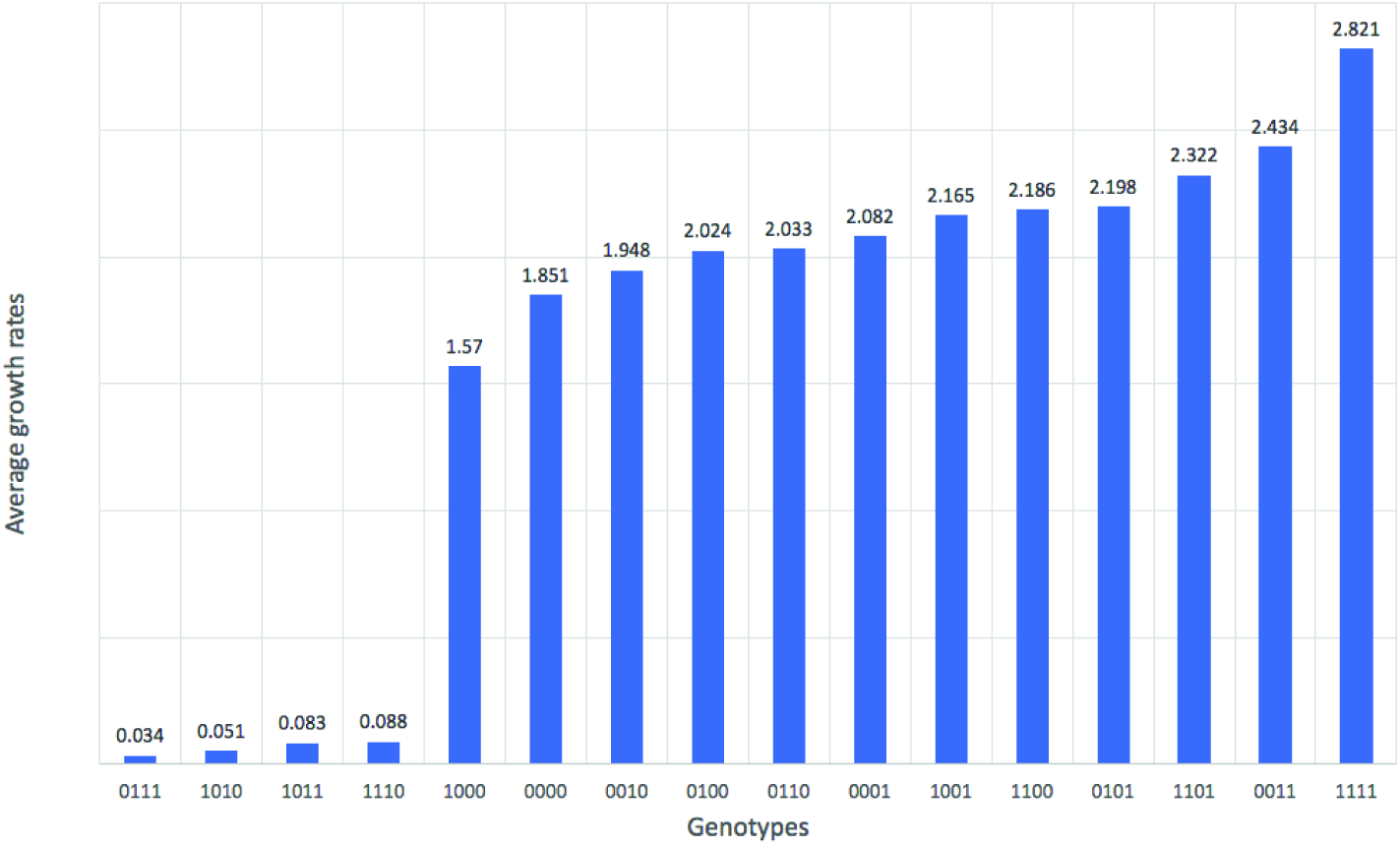
Average growth rates of 16 genotypes grown in the antibiotic AMP.

In contrast, the partial order approach, developed in this work, confirms again that the system has positive total 4-way interaction using fewer and more pronounced comparisons. To illustrate this point, we follow the notations from Theorem 2 and consider the following 8 genotypes *p*_*i*_’s and 8 genotypes *n*_*j*_’s as in Table 4. The linear form *ƒ*(*W*) is as above. The rows in Table 4 indicate a perfect matching among the two sets of genotypes. Thus, for example the first row indicates that the average growth rates associated to 1101 and 1111 are such that *w*_1101_ > *w*_1111_. Similarly, for the other rows. From these 8 comparisons of the type *w*_*p*_*i*__ > *w*_*n*_*j*__ alone, one deduces that the system has positive total 4-way interaction. Moreover, comparing with the actual average growth rates from Table 2 one can observe that the 8 differences *w*_*p*_*i*__ − *w*_*n*_*j*__ (see third column in Table 4) are bigger than the differences between the averages growth rates of the critical genotypes mentioned above. Since these differences are more significant, they provide a more reliable conclusion in the empirical setting. Finally, notice that any other perfect matching between the 16 genotypes satisfying *w*_*p*_*i*__ > *w*_*n*_*j*__ would equally well yield to the same conclusion. In summary, an advantage of the partial order approach is that it relies on fewer pairwise inequalities, which makes the approach more practical.

**Table 4.** Perfect matching of genotypes obtained from the partial fitness order according to the average growth rate.

| $i = j$ | $p_i$ | $n_j$ | $w_{p_i} - w_{n_j}$ |
| --- | --- | --- | --- |
| 1 | 1111 | 1101 | 0.499 |
| 2 | 0011 | 0001 | 0.352 |
| 3 | 0101 | 0100 | 0.174 |
| 4 | 1100 | 0010 | 0.238 |
| 5 | 1001 | 1000 | 0.595 |
| 6 | 0110 | 1110 | 1.945 |
| 7 | 0000 | 1011 | 1.768 |
| 8 | 1010 | 0111 | 0.017 |

## 7. Discussion and future directions

Understand genetic interactions from fitness measurements associated to genotypes represents a major challenge in evolutionary biology. In this work, we have focused on the case where only fitness comparisons between certain genotypes are available. This is a common assumption in practice, as there might be more uncertainty or cost involved in deducing some fitness measurements than others. Furthermore, this approach allows to exclude the measurements that have high uncertainty.

In this setting, we present a new algorithm to detect genetic interactions from partial fitness orders in an efficient way. Moreover, we derive a geometric characterization of the class of partial orders that imply interactions. This description, involving the geometry of convex polyhedral cones, provides a solid framework to develop statistical analysis of genetic interactions from partial fitness data.

Our work inspires a number of questions which remain open. First, while we are able to characterize the set of partial orders which imply interaction, our characterization is in terms of a matching in a separate graph which we can construct from our partial order. Thus, it is not immediately clear how to relate properties of a partial order to the probability that this partial order implies interaction. For instance, how does the number and type of incomparable pairs in linear orders affect whether an interaction is implied or not?

## Acknowledgments

We are grateful to Bernd Sturmfels for hosting us at the Max Planck Institute for Mathematics in the Sciences in Leipzig, where this work started. AG was partially supported by Royal Society of New Zealand through Rutherford Discovery Fellowship, contract RDF-UOO1702.

